# VoltRon: A Spatial Omics Analysis Platform for Multi-Resolution and Multi-omics Integration using Image Registration

**DOI:** 10.1101/2023.12.15.571667

**Authors:** Artür Manukyan, Ella Bahry, Emanuel Wyler, Erik Becher, Anna Pascual-Reguant, Izabela Plumbom, Hasan Onur Dikmen, Sefer Elezkurtaj, Thomas Conrad, Janine Altmüller, Anja E. Hauser, Andreas Hocke, Helena Radbruch, Deborah Schmidt, Markus Landthaler, Altuna Akalin

## Abstract

The growing number of spatial omic technologies have created a demand for computational tools capable of managing, storing, and analyzing spatial datasets with multiple modalities and spatial resolutions. Meanwhile, computer vision is becoming an integral part of processing spatial data readouts where image registration and spatial data alignment of tissue sections are essential prior to data integration. Hence, there is a need for computational platforms that analyze data across spatial datasets with diverse resolutions as well as those that manipulate and process images of microanatomical tissue structures. To this end, we have developed VoltRon, a novel R package for spatial omics analysis with a unique data structure that accommodates data readouts with many levels of spatial resolutions (i.e., multi-resolution) including regions of interest (ROIs), spots, single cells, and even subcellular entities such as molecules. To connect and integrate these spatially diverse omic profiles, VoltRon accounts for spatial organization of tissue blocks (samples), layers (sections) and assays given a multi-resolution collection of spatial data readouts. An easy-to-use computer vision toolbox, OpenCV, is fully embedded in VoltRon that allows users to both automatically and manually register spatial coordinates across adjacent layers for data transfer without the need for external software tools. VoltRon is implemented in the R programming language and is freely available at https://github.com/BIMSBbioinfo/VoltRon.

## Background

The introduction of “Spatial Transcriptomics’’ (ST)^1^ and the large-scale utilization of fluorescence in situ hybridization (FISH) techniques that allow locating mRNAs over tissue sections^2,3^ have motivated many to analyze omics data more in a spatially resolved manner. This has been followed by a rapid increase in the number of both commercially available and customized spatial omics instruments that introduced fine-tuned workflows capturing omics profiles in diverse levels of spatial resolutions^4^. The size of the area that is profiled as well as the number of genes (or features) captured by each omic profile determine the spatial resolution. Technologies that are considered to be the least spatial resolved incorporate in situ arrays, i.e. spots^5^, or user-defined segments are referred to as regions of interest (ROIs^6^). These methods often capture features across multiple cells. Whereas others with much higher spatial resolution^5,7^ may even go beyond the spatial localization of single cells, and detect the coordinates of individual mRNA molecules (i.e. subcellular resolution).

While some notable single cell analysis tools have already been adapted to store spatial localization of omic profiles and subcellular entities, a set of novel software tools and packages capable of loading, analyzing and visualizing spatial data sets are also proposed^3,8^. Seurat^9^ is one of the fundamental packages that analyze data at single cell resolution, and perhaps considered to be a gold standard for single cell analysis within the R programming community^8^. The latest version (5.0) of Seurat package introduces some number of built-in routines to load and visualize spatial data readouts, and provides a few functions to analyze data in a spatially aware manner^10^. The Giotto Suite^11,12^ has recently been developed to serve as an alternative R package where a larger set of spatially aware analysis methods are available, including neighborhood enrichment analysis and ligand-receptor pairing analysis. Unlike Seurat, Giotto suite allows storing a few different spatial units (or spatial data types) associated with each stacked layer of a single Giotto object which can be then aggregated into a single spatial unit for further downstream analysis. Squidpy^13^ is a Python based data analysis platform that extends over the already existing utilities provided by Scanpy^14^ for spatially aware analysis. SpatialData^15^ was also recently proposed to store data on spatial omics technologies using different combinations of “spatial elements’’, e.g., image, points, labels, and shapes. Squidpy and SpatialData are both currently part of the scverse ecosystem^16^. Although all these platforms are geared towards analyzing only up to supracellular (spot level) data modalities, some other software platforms to analyze regions of interest (ROIs) specifically were also introduced (SpatialDecon^17^, SpatialOmicsOverlay^18^, and StandR^19^).

Despite the existence of multiple spatially aware software platforms, image analysis is emerging as an indispensable part of spatial multimodal integration workflows ^5,20^. Spatial data sets are often accompanied by tissue images of reference that come in different flavors, e.g., H&E or DAPI^5^. In cases where adjacent or serial sections from tissue blocks are used to generate data with distinct spatial resolutions (i.e., multi-resolution) or modalities, image registration is necessary for both the alignment of morphological context across images and for synchronizing coordinate spaces of spatial data types. Currently, only a handful of omic analysis packages provide support for image-level data registration and integration. Novel tools such STUtility^21^, SpatialData and Giotto Suite allow aligning data over adjacent tissue sections by either **(i)** manually manipulating images, **(ii)** interactively picking landmark points for rigid image transformations or **(iii)** automatically align images but limited to H&E images only (STUtility). However, it may be time consuming and thus not ideal for users to manually align images, especially when dissimilar tissue staining methods such as DAPI and H&E were incorporated on each individual adjacent section. In fact, automated image registration workflows may perform better than manual approaches in these cases, specifically due to the latter being prone to human error^22^.

There remains a need for tools that analyze omic datasets with multiple spatial resolutions as well as those that incorporate more sophisticated image analysis workflows. To this end, we developed VoltRon, an R package for spatial omic analysis and data integration across serial tissue sections (**Figure 1**). VoltRon provides built-in functions to accommodate multi-resolution spatial datasets in a single R object, hence stores, analyzes and visualizes ROIs, spots, cells, and molecules, simultaneously. To accommodate such heterogeneous data, a VoltRon object is composed of multiple hierarchical data objects that are associated with tissue blocks, layers, and assays. We use this data infrastructure to seamlessly transfer and integrate data across layers of tissue blocks where assays across multiple adjacent layers are allowed to have independent sets of omic features and observations with different spatial resolutions. To integrate these features and observations across tissue layers, VoltRon utilizes multiple image alignment workflows to synchronize coordinate spaces of serial sections. We use open source computer vision libraries that include a large collection of algorithms for processing images and videos, including multiple methods for both automated and manual image registration tasks. VoltRon also acts as an end-to-end spatial analysis tool. It is possible to use a single application programming interface (API), and hence minimal number of functions, to analyze and visualize datasets with any of four spatial resolution types. In addition, VoltRon interfaces with single cell analysis tools and objects such as Seurat^9^ and SingleCellExperiment^23^ to integrate single cell and spatial datasets for spot based transcriptomics analysis. Conversion of VoltRon objects into other classes of objects used within R (Seurat) and Python (e.g., Anndata^14^) based omic analysis ecosystems are also supported.

**Figure 1:**
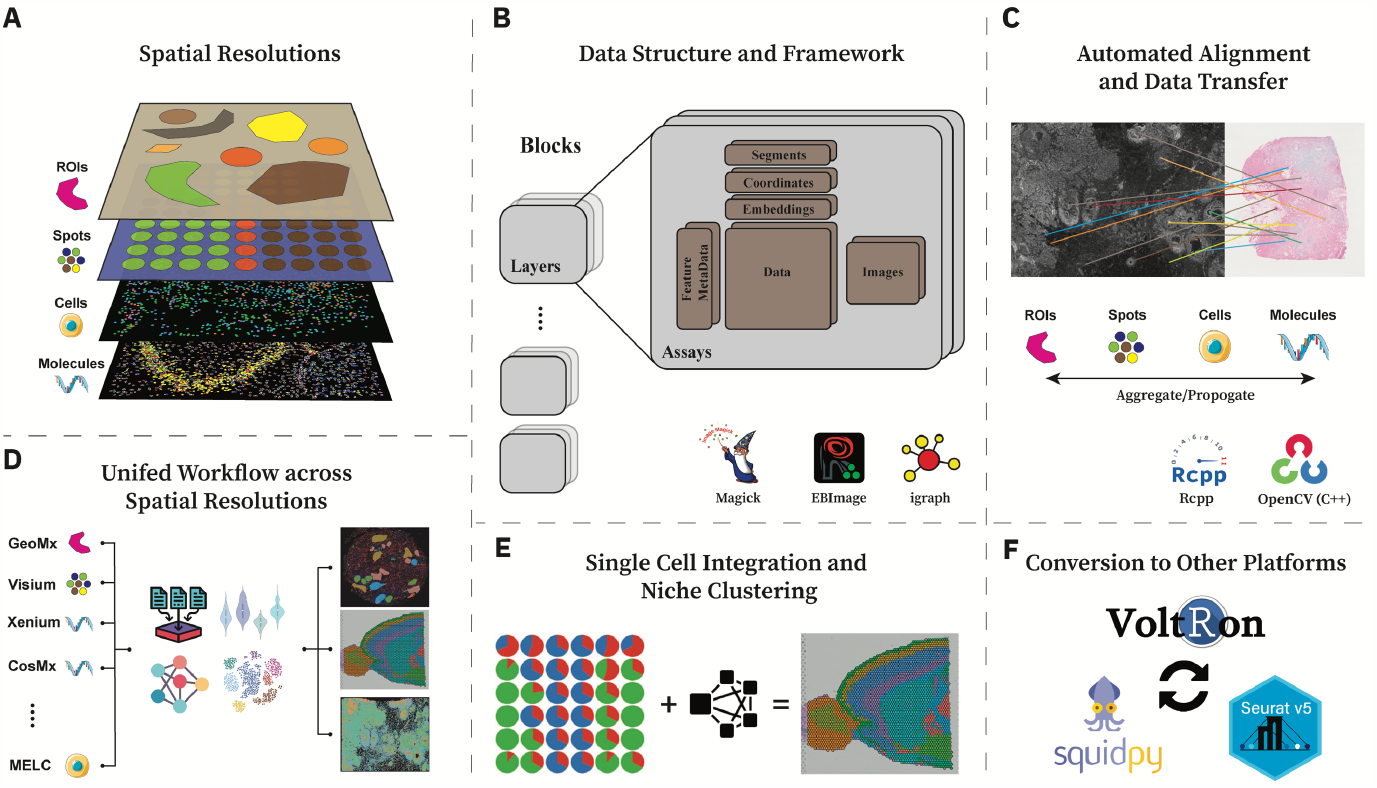
An Overview of the VoltRon package. **(A)** VoltRon can manage datasets with distinct levels of spatial resolutions: regions of interest, or ROIs (e.g., GeoMx), spots (e.g., Visium, DBIT-seq, Slide-seq^63^), cells (e.g., MELC, IMC), molecules (e.g., Xenium, CosMx). **(B)** The data structure captures the spatial organization of samples via hierarchically defined blocks, layers, and assay. **(C)** The OpenCV library is fully embedded into VoltRon for automated and manual image registration for synchronization of spatial coordinates which allows transferring data across adjacent tissue sections. **(D)** VoltRon incorporates a unified API across all four spatial data resolutions. **(E)** Integration with Seurat^9^ and SingleCellExperiment^23^ data objects provides ROI/Spot deconvolution where estimated cell type abundances could be used for niche clustering of spots. **(F)** VoltRon objects are convertible to both Seurat and Squidpy (Anndata) objects, and provide an ecosystem friendly computing environment.

Further information on the VoltRon package, including tutorials and workflows, can be accessed on https://bioinformatics.mdc-berlin.de/VoltRon. These tutorials include **(i)** integration workflows across serial/adjacent tissue sections, **(ii)** integration between spatial transcriptomics and scRNA datasets as well as **(iii)** multiple end-to-end analysis workflows for all spatial data modalities using the same API.

## Results

### VoltRon implementation and data structure

At its core, VoltRon supports downstream analysis and integration of datasets with diverse levels of spatial resolutions. One can categorize these levels as spatial data modalities with observations (*spatial points* or *spatial entities*) ranging from subcellular entities, such as molecules or transcripts, to areas over the tissue that represent large spatial segments, i.e., regions of interest (ROIs).

To accommodate different modalities and resolutions in a spatial context, VoltRon provides a hierarchical S4 data structure in R for tissue blocks, layers (sections) and assays. Thus, each block (or samples, *vrSample*) may include a number of adjacent or non-adjacent sections (or layers, *vrLayer*) where each of these layers may also contain multiple assays (*vrAssay*) that are associated to both experimentally generated (e.g. ATAC-RNA-Seq ^24^) or computationally derived (spot deconvolution results) data **(Figure 2)**. Here, assays may have independent sets of features and observations which could be subcellular entities (or molecules), cells, spots (supracellular points), and regions of interest (ROIs). VoltRon also allows defining independent sets of embeddings (for principal components and UMAP dimensions), spatial coordinates, segments, and images for each individual *vrAssay* object. Hence, VoltRon is capable of storing complex spatially-aware datasets; for example, **(i)** multiple assays within a single section such as spatial CITE-Seq^25^ and spatial ATAC-RNA-Seq^24^ that allow joint profiling of surface proteins or chromatin accessibility of spots along with gene expression, **(ii)** multiple assays with identical modalities to construct 3D spatial tissue datasets **(iii)** as well as unimodal omic data across adjacent tissue sections for more complex settings of data integration where sections were separately profiled using 10X Genomics’ Visium and Xenium instruments^5^.

**Figure 2:**
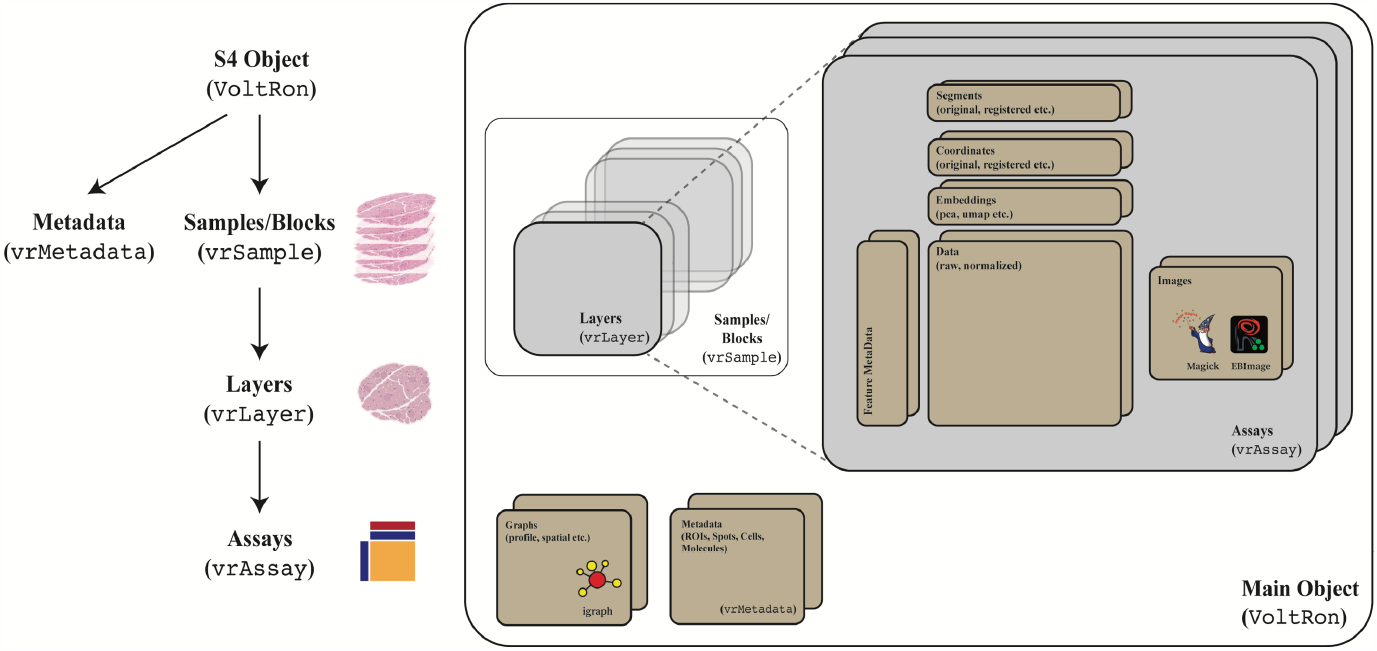
The Data Structure of VoltRon objects supports diverse spatial data modalities and spatial resolutions. A VoltRon object is composed of a set of hierarchical R S4 objects to accommodate the spatial structure of tissue sections. Each VoltRon object includes a set of *vrSample* objects representative of tissue blocks with multiple tissue sections which are themselves stored as *vrLayer* objects with each having one or more assays. Here, *vrAssay* objects store data with independent sets of feature matrices, feature metadata, images, coordinates, and other high-level information such as segments. The *vrMetadata* object stores metadata tables of each spatial resolution type.

One advantage of VoltRon is also its capability of managing multiple numbers of embeddings, images, coordinate, and segment sets in a single *vrAssay* object. Although most single cell and spatial analysis platforms store multiple lower dimensional embeddings, one may need to keep multiple coordinate (and shape data, i.e., segments) sets of the same batch of observations to account for, e.g., both unregistered and registered coordinates/images as well as points associated with different spatial resolutions of the same image.

VoltRon’s infrastructure depends on the state-of-the-art image manipulation, processing, and computer vision toolboxes. Magick^26^ and EBImage^27^ are used as the main image manipulation tools which incorporate utilities such as rotation, alteration, negation and modulation of images (**Figure 1B**). VoltRon interfaces with the open-source computer vision toolbox OpenCV (v4.7)^28^ to convert magick images into n-dimensional dense arrays which are then used to automatically capture keypoints and descriptors. OpenCV derived image registration workflows then allow matching these descriptors across images for alignment and data integration.

VoltRon provides built-in functions for importing datasets from multiple spatial omic technologies, including **(i)** imaging-based FISH assays, e.g. 10x Genomics Xenium^5^ and Nanostring CosMx^7^, **(ii)** spot-based transcriptomics assays, e.g. 10x Genomics Visium Spatial Gene Expression Platform, **(iii)** Region-specific spatial profiling technologies, e.g. Nanostring GeoMx^6^. However, one can utilize the *formVoltRon* function to build a VoltRon object from readouts of custom spatial omic technologies, e.g. DBIT-seq^29^, Slide-seq^63^, Multi Epitope Ligand Cartography (MELC)^30^ and Imaging Mass Cytometry (IMC)^31^.

VoltRon also offers end-to-end spatial omic analysis workflows for filtering, processing, clustering, and spatial association tests of spatial points or spatial entities. Although these functionalities are available in a large set of software platforms such as Seurat, Giotto Suite, Scanpy and Squidpy that are mainly geared towards single cell level data, VoltRon provides a simple programming interface that allows users to use the same set of built-in functions that react differently for each assay type **(Figure 1D)**. In addition, users can convert VoltRon objects into other publicly available data types of Seurat and Squidpy (Anndata) packages to apply additional downstream analysis workflows not available in VoltRon.

### Spatial data alignment and data transfer across layers

The most essential functionality of VoltRon is to organize spatial data readouts of distinct modalities according to the spatial context of their tissue of origin, and thus performing integration and data transfer across adjacent tissue sections. We accomplish this by incorporating a built-in mini Shiny^58^ application to align images, coordinates, segments of spatial entities of assays (**Figure 3A**). Here, coordinates and segments are transformed and aligned using the default images of their assays. Given a pair of assays and their respective images, one image is taken as a reference (destination) image and another as a query (source) image where we would like to compute a transformation matrix to achieve a perspective (homography) transformation^60^ between pixels of one image as opposed to the other (see Methods). The same transformation matrix is then used to align spatial coordinates and segments. The Shiny interface accepts a list of VoltRon objects as inputs where one image is designated as the reference (usually the central image) and the remaining images as queries. For each pair of alignment there are separate panels. Once a panel is active, all other panels associated with the same alignment task are also activated. After utilizing either the automated or manual alignment method, the app will produce additional images and slideshows that demonstrates the quality and accuracy of the alignment in the bottom-right panel.

**Figure 3:**
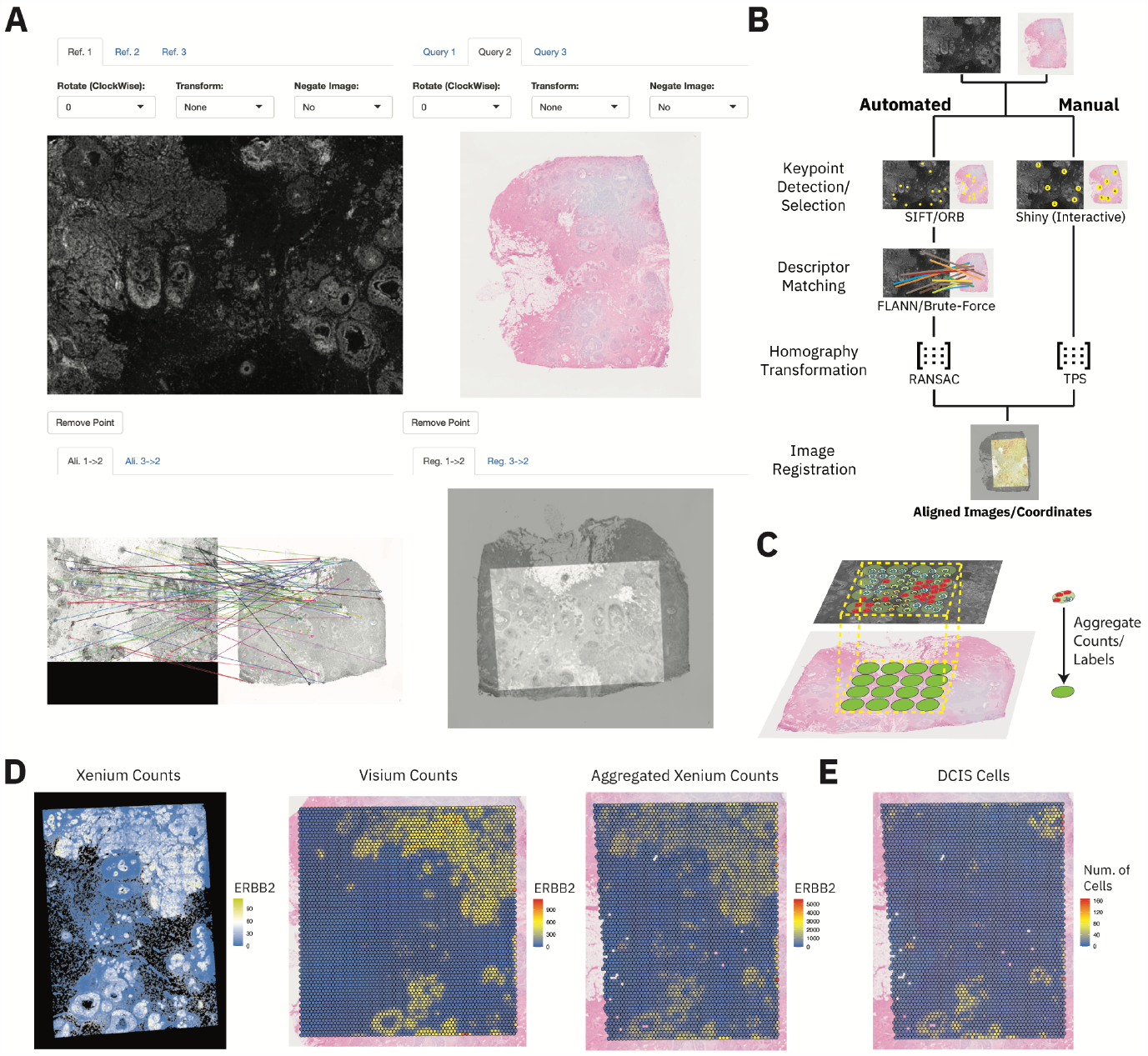
Label and data transfer across VoltRon layers (or assays) using either automated or manual image registration. **(A)** The mini Shiny application triggered by the *registerSpatialData* function. FLANN-based automated alignment is selected to align Xenium and Visium images, and the VoltRon objects with registered coordinates are returned by the application. **(B)** Image registration workflows embedded in *registerSpatialData* function using a Shiny app. Automated alignment incorporates SIFT (or ORB) for keypoint detection, matches descriptors using the FLANN (or Brute-force) method and constructs a perspective transformation matrix using RANSAC. Manual alignment also computes a perspective transformation across images using a set of interactively selected landmark points and calculates the matrix with thin plate spline (TPS) method. **(C)** An illustration of Visium spots aligning to Xenium cells after image registration. Labels and data are transferred from cells to each associated spot. **(D)** ERBB2 expression across **(i)** Xenium cells with registered coordinates, **(ii)** Visium spots and **(iii)** transferred and aggregated Xenium cells. **(E)** Transferred and aggregated DCIS sub cell types from the Xenium assay, indicating DCIS rich niches.

VoltRon streamlines the process of automatic image registration by leveraging the capabilities of OpenCV^28^ (**Figure 3B**) through a Shiny app (see Methods). OpenCV includes methods that are used to register and align multiple images by automatically detecting and matching descriptive features within each image^32,33^. The automated registration workflow of VoltRon begins by utilizing either the Scale-invariant feature transform (SIFT)^32^ or Oriented FAST and Rotated BRIEF (ORB)^33^ algorithm for keypoint detection and descriptor extraction in both the reference and query images. Subsequently, these sets of descriptors are matched using either the Fast Library for Approximate Nearest Neighbors (FLANN^64^) or a brute-force approach. The most robust matches are then selected to compute the homography transformation matrix, utilizing a Random Sample Consensus (RANSAC)^34^ iterative method. To enhance the effectiveness of the automatic registration, users are encouraged to pre-adjust the images of VoltRon objects (e.g., DAPI images) interactively — by flipping, rotating, or pixel value negating — to ensure smooth alignment. We often observe that this preliminary processing aids reducing the likelihood of failure during registration.

We demonstrate the capabilities of automated alignment between diverse spatial data modalities by registering two Xenium in situ replicates to a single Visium CytAssist assay^5^. Here, we use DAPI images from readouts of the Xenium Analyzer as query images, and the H&E images generated by the Visium protocol as the reference image. We selected the “FLANN” option (see Methods) for this registration task where SIFT is used to detect descriptors and FLANN is used to match these descriptors. We had seen great performance in the FLANN-based automated alignment when the DAPI images were negated and rotated (**Supp. Figure 1A**). A similar negation of the DAPI image was performed in tutorials provided at 10x Genomics webpage^35^. Within this supplementary guide, the authors have aligned the IF image to the associated H&E image of the same section, yet we have aligned the negated IF image to the H&E image of consecutive sections.

For cases where the automated registration approach fails, we can utilize the manual alignment method where users interactively pick locations on images of VoltRon objects. These keypoints or landmarks are expected to be pointing to similar tissue structures across both reference and query images (**Figure 3B**, see Methods). One can utilize rotation/flipping/negation operations on the images to determine the best landmarks to be used for the registration. Once landmarks are selected the Thin Plate Spline (TPS) interpolation method^36^ is incorporated to find the perspective transformation matrix which is then used to align coordinates and images of the query spatial data to the reference similar to the automated approach. It is also possible to save the selected keypoints after the alignment to reproduce the same results. We demonstrate this manual registration task by picking seven landmark points across both images where keypoints are selected from visibly clear ductal carcinoma in situ (DCIS) and invasive carcinoma niches (**Supp. Figure 1B**).

VoltRon is designed to store assays from multiple spatial data modalities within a single R object. Once aligned either using the automated and manual approach, all Xenium and Visium assays are combined in a single *vrSample* of the VoltRon object, represented as a list of distinct adjacent sections of a single block. Thus, we can transfer metadata and data features across sections. Continuous data features (gene or protein expression) can be transferred from cells to spots by aggregating cell level feature measurements to spots by detecting Xenium cells overlaid with Visium spots (**Figure 3C**). Here, we aggregate all 313 features in the Xenium assay into a new assay within the same layer of the Visium assay, thus a fourth assay is created within this tissue block where one section includes two spot level spatial assay: **(i)** one with the original Visium assay, and **(ii)** the other with aggregated Xenium cells into Visium spot level. The comparison between the original Xenium (registered) and Visium counts along with the aggregated Xenium ERBB2 counts, and aggregated DCIS cells indicate the successful alignment of two layers^5^ (**Figure 3D**). Metadata features or categorical (i.e., labels or annotations) data could also be transferred across adjacent sections. Upon label transfer, a new assay is again defined within the same layer of the Visium assay where each category of the metadata column is a feature, and the abundance of these categories are stored as raw counts. This new assay can be used to estimate the cell type abundance of the Visium spots (**Figure 3E**).

### Downstream analysis utilities for spatial omic datasets of single cell and supracellular resolution

VoltRon includes a variety of built-in functions ranging from simple image manipulations to processing, analyzing, and visualizing each of four spatial data modalities: subcellular, single cell, spot, and ROI, using the same programming interface (**Figure 1D**). Functions such as normalization, data integration and visualization react differently given each of four modalities. To illustrate some of these utilities, we first subsetted the VoltRon object interactively (similar to in **Supp Figure 3B**) for zooming on a DCIS niche in the first Xenium replicate before visualizing the localization of transcripts act as markers of myoepithelial and carcinoma cell subtypes^5^ (**Figure 4A**). VoltRon subsets assays using the background images, e.g., DAPI (or any default image within vrAssay), to zoom in and select specific niches. Here, **Figure 4A** shows this preselect niche along with the locations of two mRNA molecules serving as markers of myoepithelial cell subtypes, KRT15 and ACTA2, and two ductal carcinoma markers, TACSTD2 and CEACAM6.

**Figure 4:**
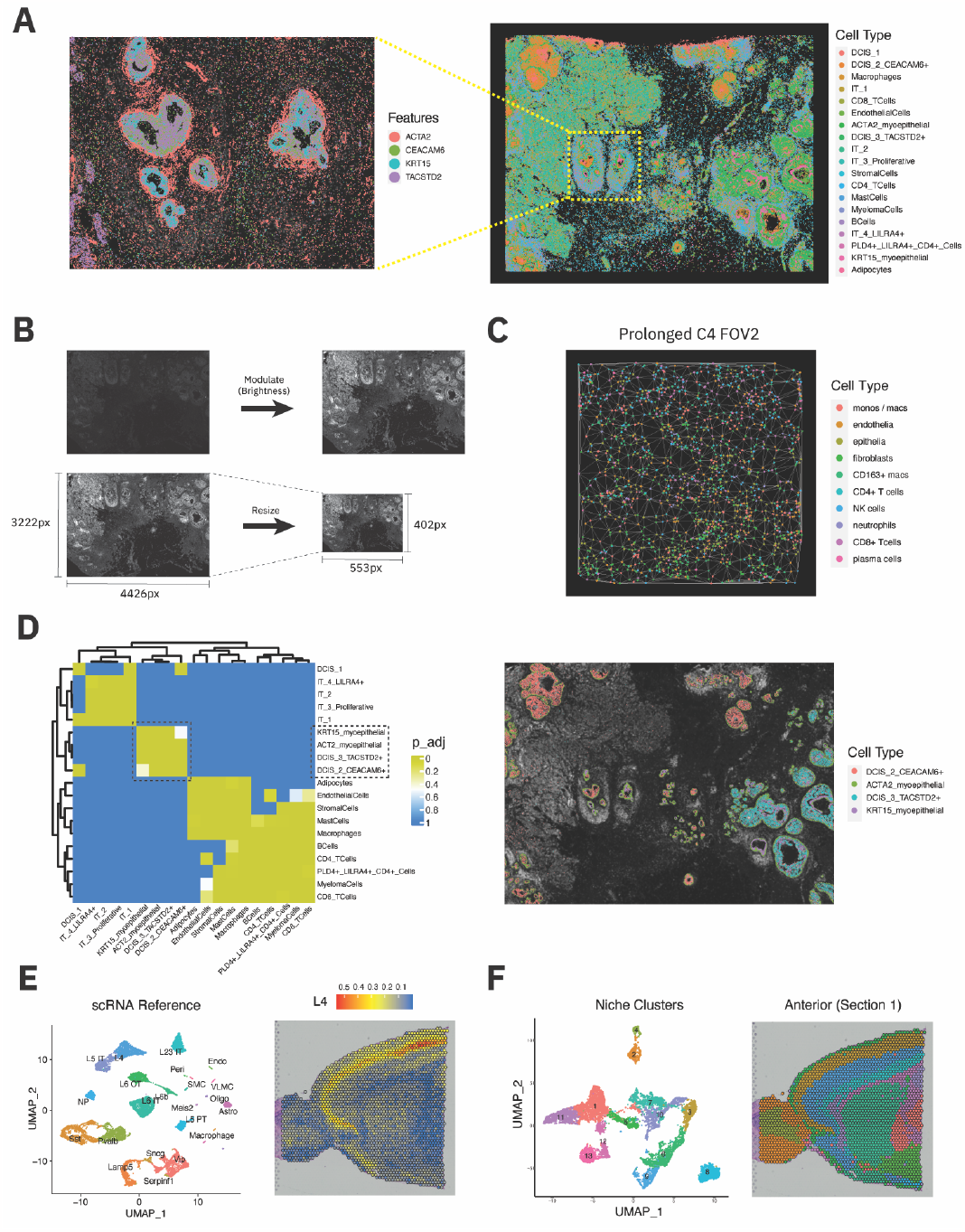
Downstream analysis utilities of VoltRon, including workflows geared towards datasets with cellular, subcellular and supracellular (spot) resolutions. **(A)** Subcellular level data visualization of Xenium transcripts for selected regions which are subsetted interactively using VoltRon (**Supp. Figure 2B**). The Xenium cells are analyzed by an end-to-end workflow ranging from filtering to clustering. **(B)** Built-in image management tools that include resizing images and modulating images for adjusting the brightness. **(C)** The Delaunay tessellation of a prolonged case 4 section (or field of view, FOV) whose edges used as a spatial neighborhood graph. **(D)** The adjusted p values of neighborhood enrichment analysis for each cell type pair, indicating a significant association of multiple cell types. **(E)** VoltRon interfaces with RCTD method^37^ for spot deconvolution of sagittal mouse brain sections (see **Supp Figure 2A**). **(F)** Cell type proportions are then used to cluster spots, i.e., niche clustering (see **Supp Figure 2B**).

Users can filter (or subset) VoltRon objects using **(i)** assays, assay types or samples, **(ii)** metadata columns, **(iii)** features, **(iv)** spatial points/entities and even **(v)** images which allows analyzing subsetting and investigating small niches. Subsetting objects are also possible using interactive Shiny apps where user-selected locations over the image are used to create new VoltRon objects (**Supp. Figure 2B**). It is possible to modulate/process the images of all VoltRon assays, and even resize, at any stage (**Figure 4B**).

VoltRon supports **(i)** depth normalization followed by log normalization, **(ii)** centered-log ratio transformation^38^ which is suitable for compositional data (see Niche Clustering within this section), **(iii)** quantile normalization for ROIs (specifically for GeoMx DSP experiments), and **(iv)** hyperbolic arcsine transformation for imaging intensity measures^39^. We analyzed a previously reported set of Multi Epitope Ligand Cartography (MELC) images^40^ derived from lung samples of a cohort of both control and COVID-19 cases stratified as acute (1-15 days, n=3), chronic (more than 15 days) and prolonged (7–15 weeks, n=3). Here, each 43 protein targets depicted by MELC were normalized using Hyperbolic arcsine transformation (with scale parameter theta=0.2) before dimensionality reduction (PCA and UMAP) and clustering. Users may incorporate internal R functions for embedding assays into lower dimensional spaces and cluster cells, or execute these capabilities in other currently existing packages by converting VoltRon objects into, e.g., Seurat or Anndata objects.

VoltRon incorporates functions to build neighborhood graphs that represent the similarity between omic profiles, in particular for clustering spots and cells. The *getProfileNeighbor* function allows building both k-nearest neighbor (kNN) and shared nearest neighbor (SNN) graphs for clustering omic profiles using the Leiden’s graph clustering algorithm^62^. VoltRon uses the igraph^41^ package to maintain both profile and spatial neighborhood graphs. For all spatial data modalities, both data and metadata features (gene expression, protein intensity, cell types etc.) can be visualized with the same set of samples; for example, gene expression and labels are visualized using *vrSpatialFeaturePlot* and *vrSpatialPlot* functions for both cellular level imaging and spot transcriptomics datasets.

Neighborhood enrichment analysis of spatial points is conducted using a permutation test^42^. VoltRon tests the association between cell type pairs (members of cell types are found near each other) and segregation (members of cell types are distant from each other) separately. For all tests, the package reports the number of each cell type and the associated p values (unadjusted and adjusted). We use the Delaunay tessellation^43^ to construct spatial graphs for each assay using *getSpatialNeighbor* function where you can also overlay these Delaunay graphs with spatial plots for each section/FOV (**Figure 4C**). Heatmap of the adjusted p values generated by the neighborhood analysis shows an association across two myoepithelial cell subtypes and DCIS cell types (**Figure 4D**).

Spot-based RNA deconvolution methods are fundamental to computational workflows for analyzing spatial assays such as the Visium^44,45^. VoltRon interfaces with Seurat and SingleCellExperiment objects to deconvolute spot assays in VoltRon objects using the RCTD algorithm^37^ (**Figure 4E, Supp. Figure 2A**). Here, we use four Visium datasets generated from two serial sagittal mouse brain sections, each divided into anterior and a matched posterior section later^61^. Spots across all these four Visium assays are deconvolved separately and the estimated cell type abundances are stored as new assays within the VoltRon layers of each original Visium assay for further downstream analysis. In these new assays, count data are now realized as compositional data matrices^38^ (column sums of data matrix add up to 1) with features being cell type (or cluster) labels which are provided by the single cell reference object used for deconvolution. Users can switch between multiple assay types of a VoltRon at any time during analysis.

Using these new assays that measure the cell type proportions per spot, one might apply a workflow to detect clusters of spots similar cell type mixtures^46^, also referred to as niche clustering. Considering that cell type proportions are categorized as compositional data, we point the readers to the existing discussions on mathematical spaces of compositional data sets^38,47^. It is important to map such compositional data measures to the Euclidean space, prior to calculating distances between data points. Thus, by normalizing the cell type abundances with centered log ratio (CLR) transformation, we can build SNN graphs using the Euclidean distances across normalized cell type proportions of spots and partition these graphs into distinct clusters showing similar patterns of cell type abundances. We can train UMAP embeddings directly from the normalized cell type abundances to visualize niche clusters, map niche clusters on spatial plots and project these abundances per cluster on heatmaps (**Figure 4F, Supp. Figure 2B**).

### Analysis of regions of interest

The real strength of VoltRon is to analyze readouts from almost all spatial technologies including segmentation (ROI)-based transcriptomics assays that capture features from large regions on tissue sections. VoltRon recognizes such readouts including ones from commercially available tools and allows users to implement a workflow similar to the ones conducted on bulk RNA-Seq datasets. To showcase VoltRon’s capabilities of analyzing spatial data modalities other than cellular and supracellular data types, we analyze morphological images and gene expression profiles provided by the readouts of the Nanostring’s GeoMx Digital Spatial Profiler (DSP) platform^6^, a high-plex spatial profiling technology which produces segmentation-based protein and RNA assays. We use user-selected segments (i.e., regions of interest, ROI) from eight tissue sections which have been collected from the same cohort investigated by MELC in the previous section^40^. These consisted of sections from control (n=2) as well as acute and prolonged COVID-19 lung (n=6) samples (Methods, **Supp Figure 3A**). We generated a total of 87 ROIs from these sections using the GeoMx DSP instrument where ROIs were categorized into distinct types during segmentation, guided by the initial staining of the tissue sections. Some of these types are fibrotic (high expression of ACTA2), immune (CD45+), epithelium (PanCK+), vessels etc. (**Supp Figure 3A**)

The library size of ROIs may depend on the type or size of the segment where some considerable number of small (filled) vessels have low counts. Hence, additional quality control measures (other than default measures generated by GeoMx DSP) could be generated by calculating the number of unique reads acquired per cell to give a glimpse of the depth of each ROI (**Figure 5A**). By visualizing the “CountPerNuclei”, we can see that most of these small filled vessels have in fact considerably high transcript count per cells, but five ROIs have extremely low counts. Correlating the per cell counts with the sequencing saturation, we can see that the percentage of copies is considerably low (∼25%) for these five ROIs (**Figure 5A**). Hence, we may filter out ROIs whose count per nucleus/cells are less than 500. VoltRon includes multiple preprocessing options where **(i)** ROIs could be filtered for given metadata features (e.g. CountPerNuclei > 500), **(ii)** genes with low maximum count could be removed and/or **(iii)** The 3rd quartile normalization can be applied to these reads according to the manufacturer’s specifications^48^.

**Figure 5:**
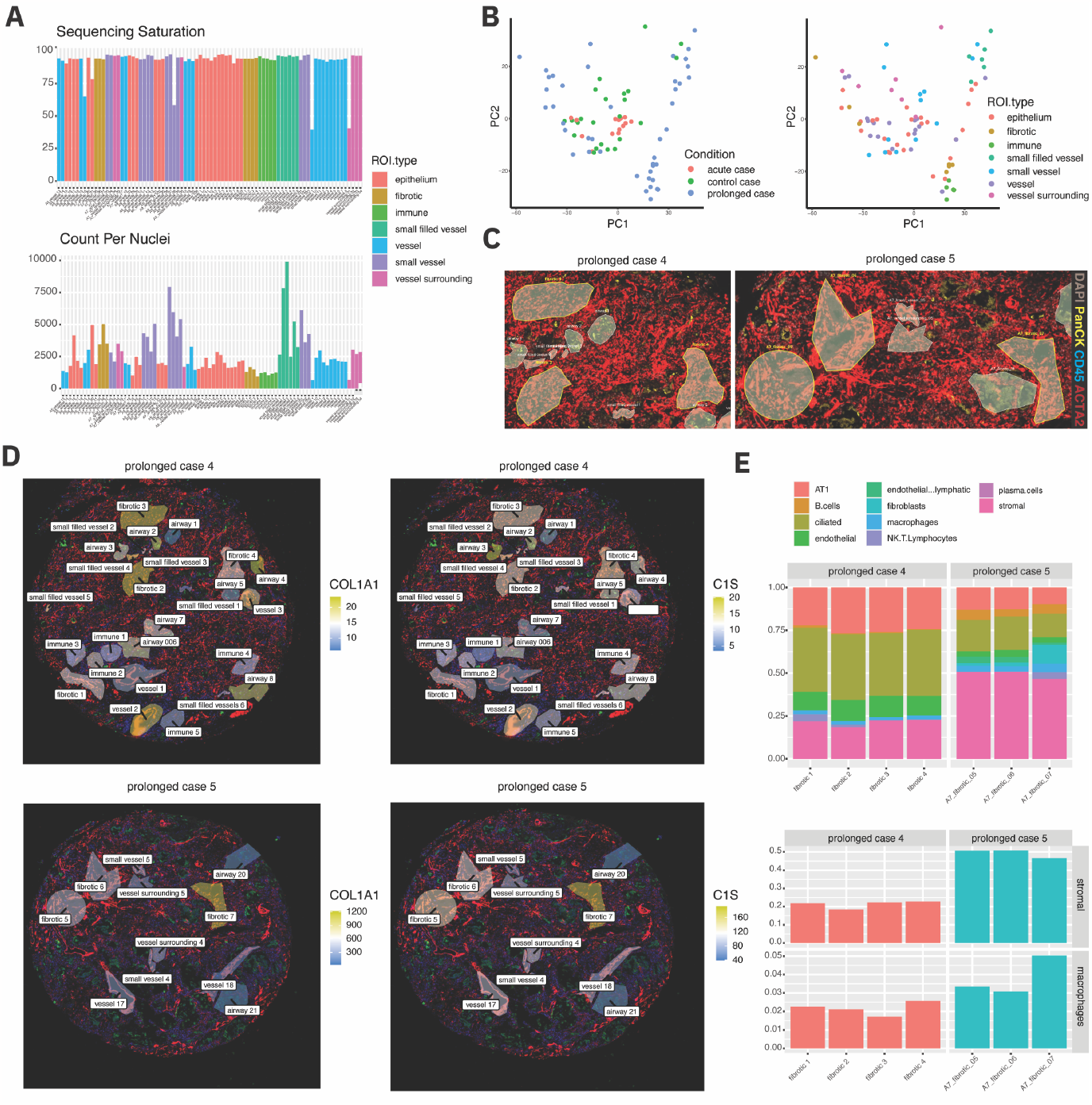
Downstream analysis of GeoMx DSP datasets: visualization, filtering, normalization, differential expression analysis and ROI deconvolution. **(A)** Bar plots of GeoMx ROIs visualize the sequencing saturation (SS) and the count per nuclei (CPN) measures for quality control. Five ROIs show low SS and CPN, thus filtered out and removed. **(B)** PCA embedding of conditions and ROI types showing heterogeneity across ROIs, specifically across prolonged COVID-19 cases. **(C)** Zoomed in microscopic images (available in GeoMx DSP instrument) of two prolonged cases, highlighting (yellow border) three of the fibrotic regions from each prolonged case. DAPI (grey), PanCK (yellow), CD45 (blue) and ACTA2 (red) **(D)** Visualization of the ROI segments of two prolonged lung sections with ROI names and normalized expression of fibrosis markers COL1A1 and C1S. **(E)** ROI deconvolution of fibrotic ROIs showing differentially abundant cell types. The fibrotic regions of prolonged case 5 show increased abundance of smooth muscle cells and macrophages.

Interactive subsetting utilities of VoltRon could be used for multiple purposes beyond selecting niches within individual tissue sections. The collection of eight lung tissue sections is initially put into a single GeoMX slide (see Methods) and could be imported into VoltRon as one object, which then are separated into several VoltRon objects using the built-in mini Shiny app (**Supp. Figure 3B**). We derive multiple subsets from the GeoMx slide image and divide the GeoMx readouts into multiple ROI-based readouts.

To investigate the heterogeneity of ROIs across these eight lung sections, we first dimensionally reduced the transcriptomic profiles of all ROIs (**Figure 5B**). We selected the 3000 genes with highest variability whose rank were calculated using the variance stabilizing transformation^49,50^ and then reduced the dimensionality of these genes using PCA. Embedding plots of the first two principal components show a great heterogeneity across ROIs of two prolonged tissue sections (case 4 and 5) where we also see a clear clustering of fibrotic and immune regions. This might indicate a considerable difference across fibrotic regions of prolonged lung sections (**Figure 5C**). To this end, we visualized and compared the spatial expression patterns of fibrosis markers such as COL1A1 and C1S for ROIs (**Figure 5D**). There seems to be an increase in the expression of both markers where the increase in COL1A1 expression is more pronounced. Visualizing the features on spatial images indicate that the among three fibrotic regions of the prolonged case 5, third and the remote region with ID “fibrotic 7’’ show higher expression of COL1A1.

Building on our observations in the embedding plots of ROIs, we can detect differentially expressed genes across ROI types (fibrotic, immune, epithelium etc.). VoltRon interfaces with DESeq2^51^ to provide support for identifying ROI types that have distinct transcriptional profiles. Correlating with the PCA embedding plot, we see two sets of fibrotic regions with distinct gene expression patterns (**Supp Figure 3C**). Subsetting the VoltRon object for these seven fibrotic regions, and testing for the differential expression between two prolonged samples (case 4 and 5), we see that an increased expression of COL1A1 and C1S along with CD68, a marker of macrophages and FN1, a profibrotic marker expressed by macrophages. We then asked if the increase in these markers could be validated by estimating the abundance of macrophages in fibrotic ROIs of prolonged case 5. By interfacing with the MuSiC deconvolution method^52^, we observed an increase in the macrophages of these fibrotic regions (**Figure 5E, Supp Figure 3D**). VoltRon requires scRNA datasets as reference, either as Seurat or SingleCellExperiment objects, for ROI or Spot deconvolution. We observed an increase in the abundance of cells that are scRNA clusters labeled as macrophages and stromal^40^ (associated with both vascular and airway smooth muscle cells). VoltRon incorporates a single function for deconvoluting both spot and ROI assays where the function runs either RCTD or MuSiC methods depending on the detected assay type.

## Discussion and Conclusion

Here we describe VoltRon as a computational platform for analyzing and integrating spatial omic readouts with diverse modalities and spatial resolutions. VoltRon provides support for defining multiple chunks (i.e., blocks/samples) of adjacent and serial tissue sections (and their associated spatial data readouts) to transfer data or metadata features across layers upon image registration. Both automated and manual alignment of tissue images are available with easy-to-use interactive Shiny interfaces where users can manipulate images or choose landmark points prior to the alignment. We also utilize a powerful opensource computer vision toolbox, OpenCV, to embed automated image registration routines, and to align images and coordinates in a quick manner. These registration workflows generate homography transformations across images which are then used to synchronize coordinates of spatial points across multiple sections into a single coordinate system. We showcased this utility by aligning and transferring data across CytAssist Visium and Xenium in situ assays whose readouts provide diverse microscopy images (DAPI vs H&E) that a manual image registration approach may be prone to human error. VoltRon does not require any additional step for configuring and compiling OpenCV, C++ or other external software/dependencies. VoltRon can easily be installed on multiple operating systems.

To achieve a more feasible integration of spatial entities across tissue layers, VoltRon offers a novel data structure to hierarchically define tissue blocks, layers, and assays which preserves the spatial organization of the data of origin. A VoltRon object can store multiple layers within a single tissue block as well as multiple assays within a single layer, allowing to identify both uni-modal and multi-modal data of tissue sections across and within distinct samples/blocks. Moreover, all assays within a VoltRon object can accommodate independent sets of observations with any spatial resolution (ROIs, spots, cells, and molecules) and diverse feature types (RNA, ATAC, protein or custom features, e.g., cell type proportions for niche clustering, see **Figure 4F**). This includes multiple images, spatial coordinate matrices and segments per assay, hence both original and unregistered coordinates of ROIs, cell segmentations and centroids could be stored within a single assay. Thus, this spatial data structure allows managing datasets of multiple data types within a single R object and simplifies the programming experience such as in the use case where cellular Xenium assays were integrated to the supracellular Visium assay. We further demonstrated the flexibility of VoltRon by analyzing datasets from other spatial omic technologies like MELC and GeoMx. Unlike currently available software platforms, VoltRon is capable of accommodating and analyzing a larger selection of spatially resolved datasets, including those with ROI resolution. In Table 1, we compare some of the fundamental utilities of VoltRon with the ones from other software platforms that analyze spatial omic datasets.

**Table 1:**
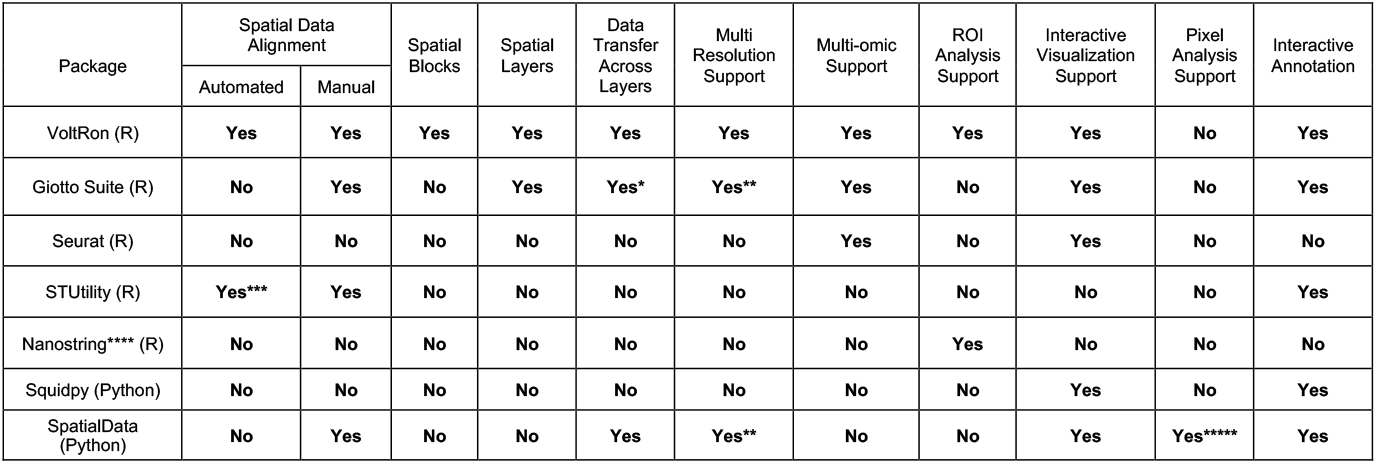
The comparison between essential utilities of VoltRon and similar functionalities available across other spatial omic analysis platforms. (*) Data transfer is achieved by aggregating layers. (**) Multiple resolutions cannot be stored within a single SpatialData object, but it is available in Giotto Suite (only up to supracellular/spot assays). (***) only for H&E images, by detecting tissue borders. (****) This includes a large collection of computational tools, GeoMxTools^53^, SpatialDecon^17^, SpatialOmicsOverlay^18^, provided by Nanostring Technologies Inc. (*****) Tiles or pixels are not spatial elements but instead are used to train Deep learning models^54^. (*,**) SpatialData and Giotto Suite are both currently under development, and these utilities are subject to change in the future.

As for potential improvements to the VoltRon platform, we will offer more comprehensive data integration workflows across adjacent/serial tissue sections, including 3D spatial data reconstruction. We will also define additional spatial resolution types such as pixels (or tiles). Consequently, convolutional autoencoder (CAE) based embeddings could be trained on pixels or tiles of images along with single cell gene expression profiles, across assays of either the same or adjacent sections, for multi-modal clustering or inference. Pixels/tiles could also be used to analyze the spatial context of multiplex tissue imaging datasets as previously reported^55^.

Interactive visualization of spatial omics data, and more importantly, co-visualizing embedding and spatial coordinates of spatial entities is crucial for the interpretation of downstream analysis results. Currently, VoltRon provides support for interactive spatial data visualization of only the centroids of cells and spots via Vitessce^56^, a novel integrative data visualization framework that could be used in R packages and Python environments. Future versions of VoltRon will offer more routines derived from currently available packages such as the vitessceR, an R package wrapper for the Vitessce platform. Although it is currently possible to convert VoltRon objects into Seurat and Anndata objects, more built-in conversion functions will be implemented to convert VoltRon object into Giotto objects, and save as other formats (Zarr^15^ and tileDB^57^) currently popular within the spatial and single cell analysis ecosystems.

## Methods

### GeoMX™ Digital Spatial Profiling of Lung Tissue Sections

Control and COVID-19 lung biopsy samples were collected postmortem (within 3 to 82 hours), fixated, and embedded in paraffin as previously described^40^. The FFPE samples were stored at room temperature for 3 years after collection. Cores of 2mm diameter were punched out of FFPE samples and incorporated into one tissue multi array (TMA), and then sliced into 5 μm sections. Deparaffinization, antigen retrieval and post-fixation were performed for GeoMx DSP workflow according to the manufacturer’s instructions. Slides were incubated overnight with the Whole Transcriptome Atlas RNA Hs probe set, followed by stringent washes. Slides were stained with fluorescently labeled antibodies against CYTO 13 (GMX-MORPH-NUC-12, NanoString) for DNA staining, PanCK (NBP2-33200, Novus), CD45 (NBP2-34528, Novus) as well as alpha smooth muscle actin (ACTA2, ab202296, abcam). Following image registration and barcode collection on the GeoMx instrument, collected barcodes were amplified with 18 PCR cycles and sequenced on a NovaSeq 6000 instrument to a depth of 100 reads per μm2. GeoMx NGS Pipeline was used to process the FASTQ files and generate DCC files.

### Spatial Data Alignment

We use the *registerSpatialData* function in the VoltRon package to execute a mini Shiny^58^ application for image registration. This interface allows rotating, flipping and negating images which are available using the magick^26^ package in R. OpenCV^28^ implementation in VoltRon is fully embedded using Rcpp^59^. Upon installation of the package, OpenCV (v4.7) is automatically imported and all C++ source functions are compiled in Mac, Windows, and Linux-based operating systems.

### Automated Image Registration

There are two available automated registration workflows in the Shiny application (*registerSpatialData* function) that incorporate either the SIFT or ORB methods for detecting keypoints: **(i)** For SIFT^32^, we performed keypoint detection and feature descriptor extraction on both reference (destination) and query (source) images, using default arguments. The descriptor matching was accomplished using the Fast Library for Approximate Nearest Neighbors (FLANN^64^, *cv::FlannBasedMatcher*) with 5 parallel KD-Trees (*trees=5*) and set to perform 50 recursive checks (*checks = 50*). Key points were then filtered using Lowe’s ratio test with a threshold of 0.8. **(ii)** In the ORB^33^ method, after keypoint detection and descriptor extraction, a BruteForce-Hamming matcher (*cv::DescriptorMatcher* with type *BruteForce-Hamming*) was used to determine the distances between feature descriptors, selecting the best P% (*GOOD_MATCH_PERCENT)* from a number of keypoint descriptor pairs (MAX_FEATURES). For both SIFT and ORB approaches, these selected pairs were used to compute the homography matrix for transforming from the source to the destination image space, employing an iterative RANSAC^34^ process with a threshold of 5 to ensure robustness against outliers. An example of this brute force approach between two Visium assays can be found at https://bioinformatics.mdc-berlin.de/VoltRon/registration.html#Alignment_of_DLPFC_Visium.

### Manual Image Registration

Manual registration option in the Shiny application (*registerSpatialData* function) allows selecting landmark points (or keypoints) across the reference and query images by interactively selecting locations over images. At least three landmark points should be chosen from both images of each pair. Users can deselect points by clicking on “Remove Points” buttons below each image. Thin plate spline interpolation method is used to calculate the perspective transformation matrix^36^.

## Data Availability

Xenium replicates and the associated Visium CytAssist data can be accessed at https://www.10xgenomics.com/products/xenium-in-situ/preview-dataset-human-breast. Visium dataset of mouse brain serial sections from Sagittal-anterior and-posterior regions can also be found at https://www.10xgenomics.com/resources/datasets. MELC feature matrices and spatial coordinates as well as microscopy images are publicly available in the Zenodo repository under DOI: 10.5281/zenodo.744749. Processed GeoMx DSP data can be accessed from the ROI analysis tutorial of VoltRon website (https://bioinformatics.mdc-berlin.de/VoltRon/roianalysis.html).

## Code Availability

The VoltRon package (R) is freely available at https://github.com/BIMSBbioinfo/VoltRon. The scripts for generating results in this manuscript are largely available under “Explore” section of the VoltRon website: https://bioinformatics.mdc-berlin.de/VoltRon/tutorials.html

## Acknowledgements

The authors are most grateful to the donors and their relatives for consenting to autopsy and subsequent research, which were facilitated by the Biobank of the Department of Neuropathology, Charité– Universitätsmedizin Berlin and the German National Autopsy Network (NATON - 01KX2121). The study was approved by the local ethic committee at Charité (approval numbers: EA1/144/13, EA2/066/20, EA1/075/19) and the Charité-BIH COVID-19 research board. All autopsies were performed at the Department of Neuropathology and the Institute of Pathology, Charité–Universitätsmedizin Berlin. Autopsy consent was obtained from the patients or their families.

## Author Contributions

AM and AA wrote the manuscript, with contributions from EBa, EW and EBe. EBa and DS have developed the automated image alignment workflows. EBe, APR and HR have prepared lung tissue sections prior to GeoMx DSP run and selected ROIs. HOD and SE have collected the biopsy samples of the GeoMx tissue sections. IP prepared the slides, initiated GeoMx DSP run and prepared the sequencing library. HR, TC and JA have supervised the GeoMx slide preparation and processing. AM, EW, EBe, and APR have analyzed the GeoMx and MELC data. AA, ML, DS, HR, AH and AEH have supervised the entire project. All authors approved the manuscript.

---

**Supp. Figure 1:**
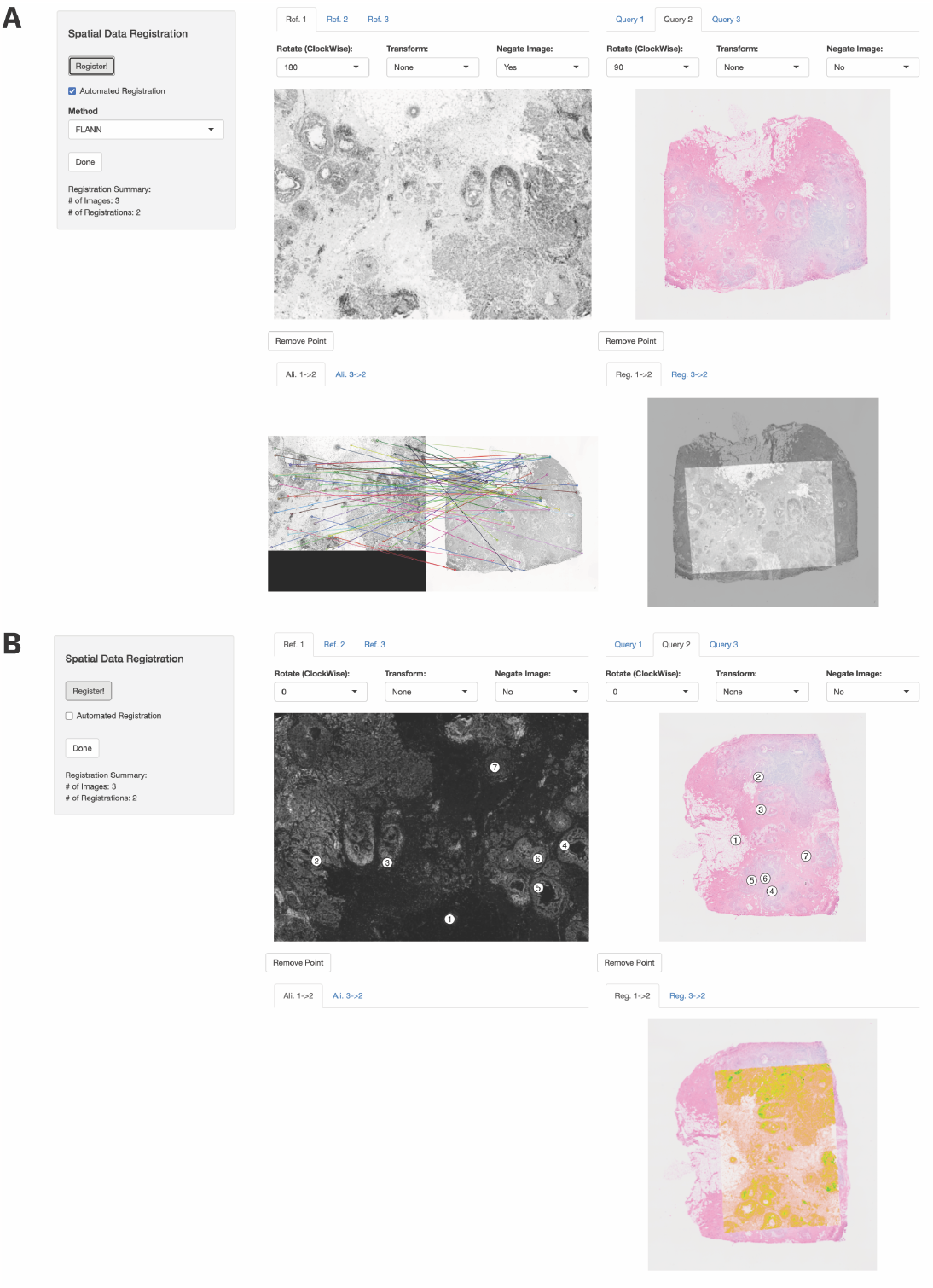
The Shiny app based interactive image alignment and registration interface. The *registerSpatialData* function accepts a list of VoltRon objects and returns (upon pushing the “Done” button) the same list with each VoltRon object with registered coordinates and, if exists, segments. The interface is divided into four panels where the first two on top visualizes each pair of adjacent sections for keypoints selection and detection for pairwise descriptor matching and alignment. The last two on bottom panels show the matched keypoints and the registered image overlayed on the reference image per each selected pair. All images can be rotated, flipped (both horizontally and vertically) and negated. **(A)** The automated alignment mode is triggered by clicking on the “Automated Registration” tick box. There are two automated alignment workflows, named FLANN and BRUTE-FORCE (see Methods). An additional set of parameters are requested upon selecting the BRUTE-FORCE option. **(B)** For manual registration, keypoints (or landmark points) are selected manually in order where numeric indices indicate matching keypoints across two images. Selected keypoints can be removed using the “Remove Point” button.

**Supp. Figure 2:**
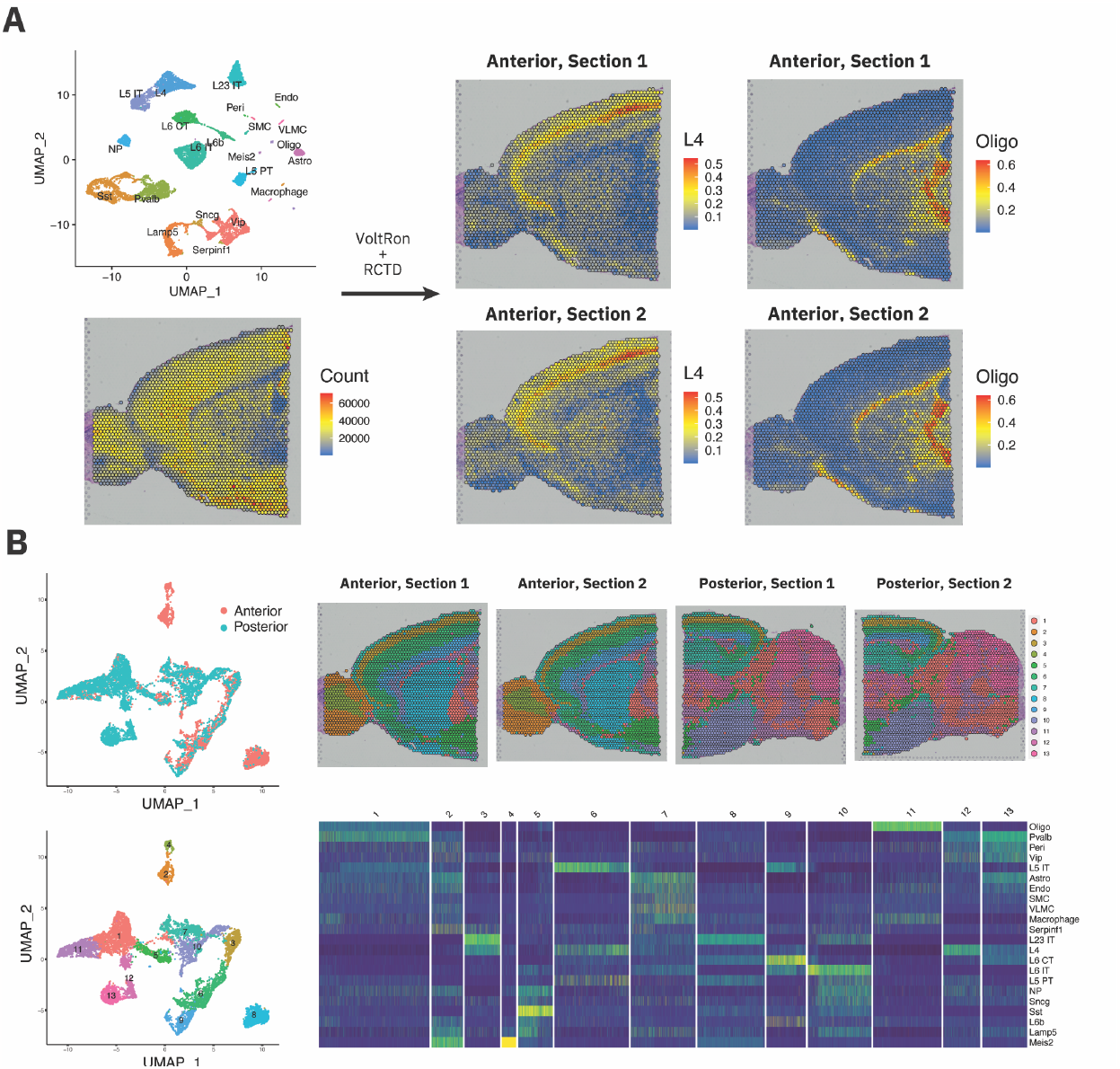
Niche Clustering of sagittal mouse brain sections via Spot Deconvolution **(A)** VoltRon interfaces with RCTD method^37^ and Seurat objects to deconvolute Visium spots, and to build new cell type abundance assays. **(B)** Cell type proportions are used to cluster spots into groups with similar cell type mixtures (i.e., niche clustering) and calculate UMAP directly from normalized cell type abundance data. Heatmap shows distinct cell type mixture patterns across niche clusters.

**Supp. Figure 3:**
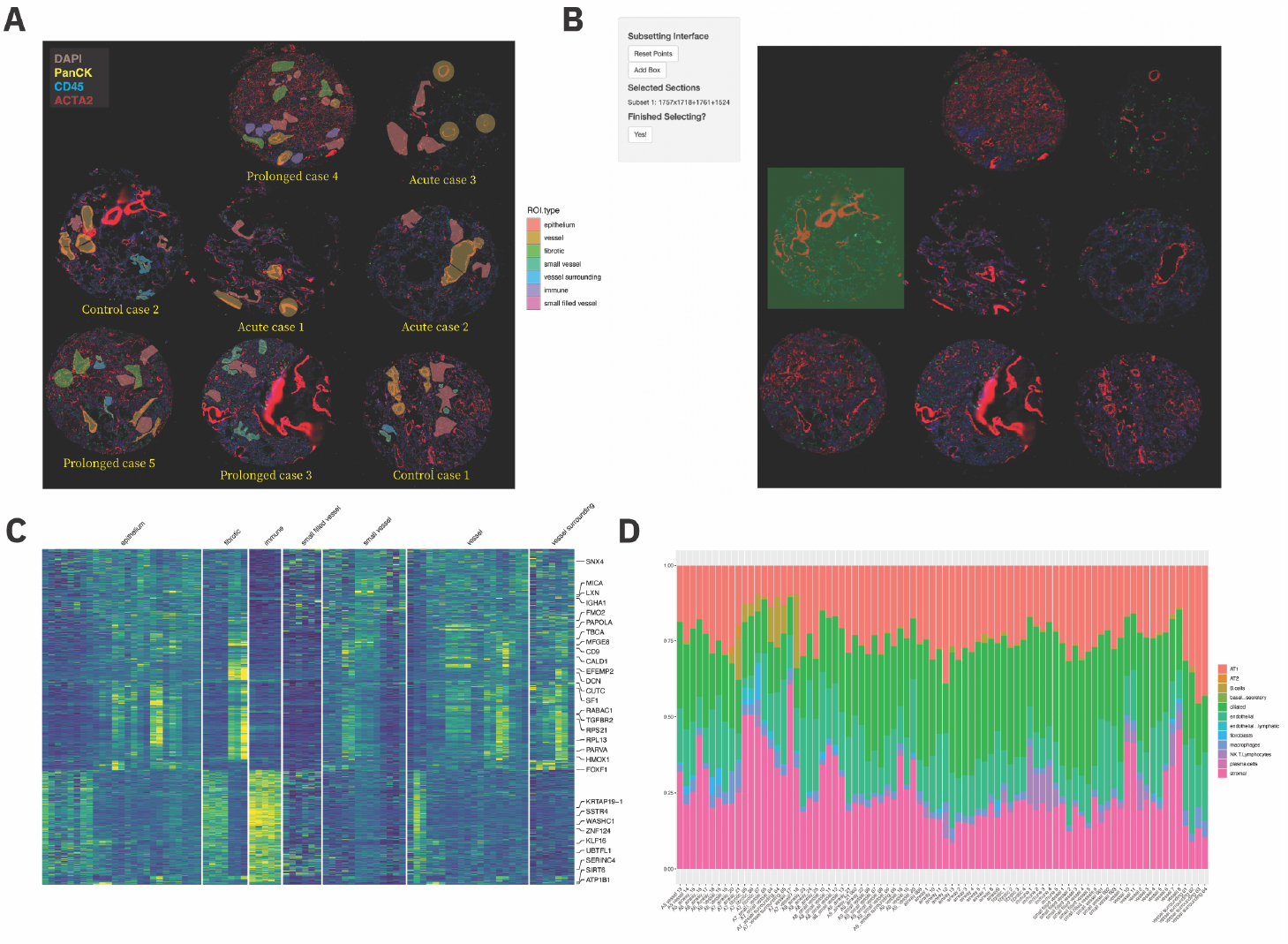
Downstream analysis utilities for GeoMx and region of interest (ROI) based spatial assays. **(A)** Spatial visualization of 87 ROIs with respect to ROI types: epithelium, fibrotic, immune, (small/small-filled) vessel, vessel surrounding. Here, there are eight adult human lung tissue sections that are categorized as the control case as well as acute and prolonged COVID-19 cases. **(B)** Built-in support for subsetting spatial datasets is used to subset tissue sections from a single GeoMx scan area. A mini shiny app is triggered using the generic R *subset* function with the “interactive=TRUE” option. Multiple subsets can be defined at a time. **(C)** A heatmap of differentially expressed genes (|logFC| > 1 and p.adj < 0.05) across ROI types (one vs all per each ROI type), indicating two distinct groups of fibrotic regions. DE genes are found by interfacing with the DESeq2^51^ package through VoltRon. **(D)** Proportion plot of ROI deconvolution results using a single cell reference dataset. There are increased abundances of stromal (ACTA2+) cells in some fibrotic regions and increased NK and T cells abundance in immune regions.

